# Membrane systems for lipid secretion in the hairy roots of *Lithospermum erythrorhizon* observed by electron microscopy

**DOI:** 10.64898/2025.12.14.694173

**Authors:** Shingo Kiyoto, Takuji Ichino, Tatsuya Awano, Kazufumi Yazaki

**Affiliations:** Laboratory of Tree Cell Biology, Division of Forest and Biomaterials Science, Graduate School of Agriculture, Kyoto University, Sakyo-ku, Kyoto, Kyoto, 606-8502, Japan; Laboratory of Medical Cell Biology, Kobe Pharmaceutical University, Kobe, Hyogo, 658-8558, Japan; Laboratory of Plant Gene Expression, Research Institute for Sustainable Humanosphere (RISH), Kyoto University, Uji, Kyoto, 611-0011, Japan

**Keywords:** Freeze-substitution, Hairy root, Lipid secretion, *Lithospermum erythrorhizon*, Multivesicular body, Shikonin, Transmission electron microscopy

## Abstract

Many plant-derived lipophilic compounds benefit human life. For example, shikonin derivatives—lipophilic red pigments from the medicinal plant *Lithospermum erythrorhizon*—exhibit wound-healing and antimicrobial effects. These compounds are excreted from root epidermal cells and accumulate in the root bark. Our understanding of how lipophilic compounds are transported in plant cells remains limited, and *L. erythrorhizon* hairy roots serve as a good model for studying the lipid secretion. To investigate this, the shikonin secretion was induced in cultured hairy roots, which were then fixed via chemical fixation or high-pressure freezing followed by freeze-substitution and observed by transmission electron microscopy. Electron micrographs from chemically fixed samples showed multivesicular bodies (MVBs) surrounding electron-dense bodies—presumed to contain shikonin derivatives—and fusing with the plasma membrane. Similarly, freeze-substituted samples revealed MVBs fused with the plasma membrane. Additionally, some MVBs contacted plastids containing enzymes for shikonin precursor biosynthesis. These findings suggest MVBs play key roles in both synthesis and secretion of shikonin derivatives.

**Highlight:** This study reveals that multivesicular bodies mediate both synthesis and secretion of shikonin derivatives in the hairy roots of *Lithospermum erythrorhizon*.

## Introduction

Plants produce lipophilic secondary metabolites to adapt to their environment. Lipophilic polymers such as wax, cutin and suberin have been intensively studied. In contrast, in plant secondary metabolism many lipophilic low-molecular-weight compounds also play important roles in defense against oxidative stress, pathogens, and herbivory. These compounds, including monoterpenes, alkaloids, and prenylated phenolics such as flavonoids and coumarins, have also been utilized by human life as medicinal resources, aromatics, and natural dyes. Notably, many of those lipophilic secondary metabolites are secreted out of the cells and accumulated in the apoplastic spaces (Ichino and Yazaki, 2022). Because these metabolites are hydrophobic, they require specialized mechanisms for transport within and between plant cells, which exist in a hydrophilic environment. The transport of these compounds is essential for their proper function, making it crucial to understand their transport mechanisms for plant bioengineering applications.

Recent advances in plant cell biology and molecular genetics have elucidated two main pathways involved in the trafficking of lipophilic secondary metabolites: the ATP-binding cassette (ABC) transporter-mediated pathway, particularly G-type (ABCG) transporters (Li *et al*., 2022), and the vesicle-mediated pathway (Ichino and Yazaki, 2022). ABCG transporters facilitate the apoplastic accumulation of various compounds, including waxes, cutin, and secondary metabolites (Pighin *et al*., 2004; McFarlane, 2010; Fu *et al*., 2024). The vesicle-mediated pathway plays a significant role in the intracellular movement and the secretion of lipophilic metabolites within membrane-bound vesicles (McFarlane, 2014; Ichino *et al*., 2020). Despite these insights, detailed ultrastructural observations of intracellular and intercellular transport of these compounds remain limited, highlighting the need for further research.

Here, shikonin derivatives (Fig. 1), which are lipophilic red naphthoquinone pigments secreted from the medicinal plant *Lithospermum erythrorhizon* Sieb. et Zucc. (Boraginaceae), exhibit anticancer, anti-inflammatory, wound-healing, and other various beneficial properties (Yazaki *et al*., 2017; Guo *et al*., 2019). *L. erythrorhizon* provides a good model for investigating membrane dynamics associated with lipid metabolism because the cultivation methods for callus and hairy roots are well-established (Tabata *et al*., 1974; Shimomura *et al*., 1991), and the production of shikonin derivatives can be robustly induced by the shikonin-production medium M9 (Fujita *et al*., 1981), and the shikonin induction pattern can be easily confirmed by red-colored culture medium and the plant tissue or roots. Hairy roots of this plant are a better model than callus tissues because hairy roots have histological structures more similar to those of native plants and an efficient transformation method by *Rhizobium radiobacter* strain A13 was established in the recent study (Tatsumi *et al*., 2020). In addition, hairy roots are suitable for freeze-substitution (FS) because they are small in thickness and therefore, are easily vitrified for small thermal capacity.

**Fig. 1.**
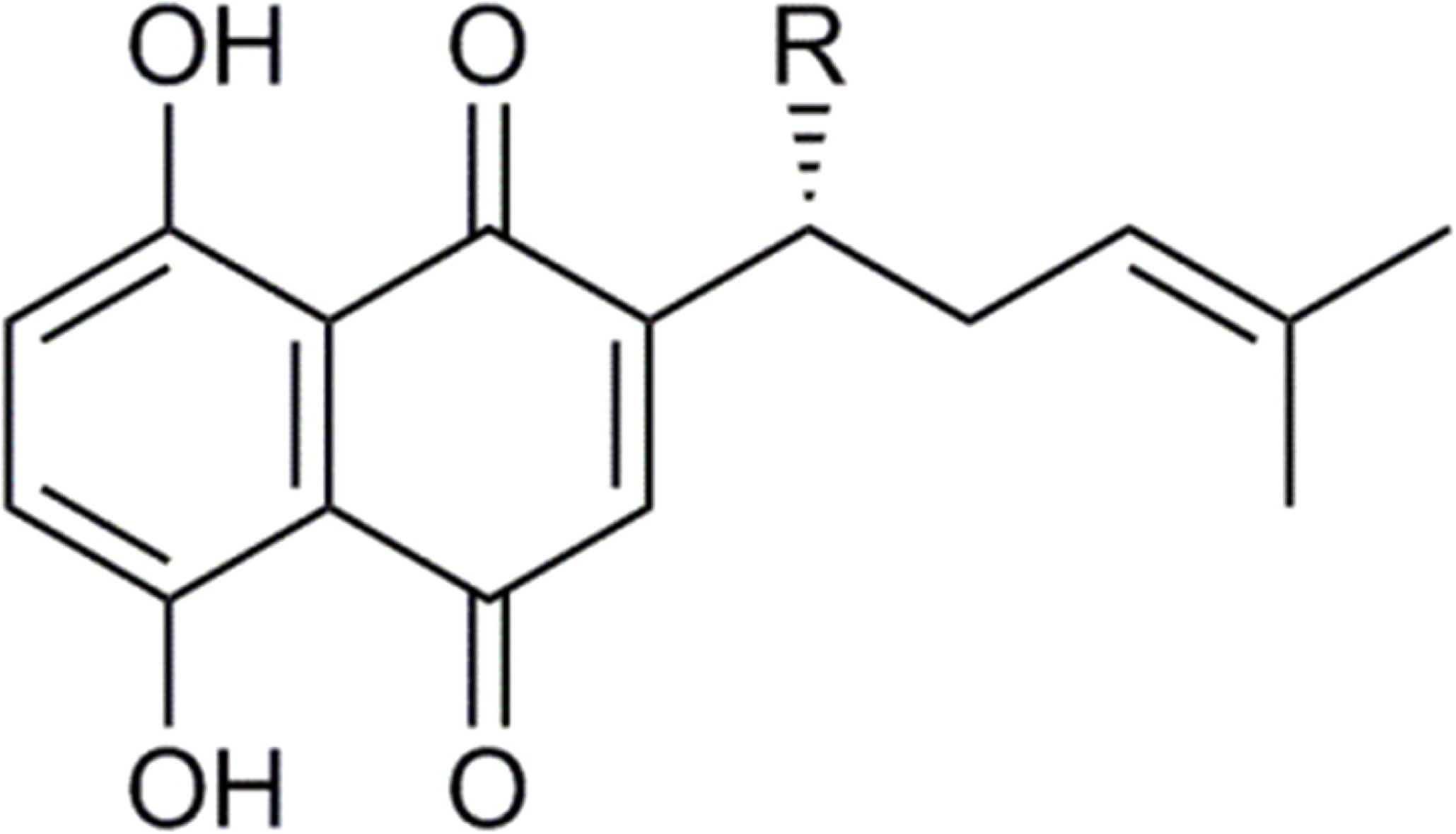
Chemical structures of shikonin derivatives produced by *L. erythrorhizon*. R reveals the fatty acid residues.

Transmission electron microscopy (TEM) is one of the popular methods for ultrastructural observation of living tissues. It requires a fixation process to prevent autolysis, decay, shrinkage, and displacement and to abolish membrane function for resin embedment. We established a chemical fixation (CF) protocol for visualize lipophilic compounds and subcellular structures (Kiyoto *et al*., 2022). In the research, the invagination of plasma membrane was observed in the cells secreting shikonin derivatives. However, such membrane deformation may be an artifact caused by CF, such as bacterial mesosomes (Nanninga, 1968;1969;1971; Ebersold, 1981). CF at room temperature can cause membrane deformation because of the slow penetration of osmium tetroxide (Hagström and Bahr, 1960) and can lead to incorrect interpretation.

FS after rapid freezing or high-pressure freezing is a solution to minimize deformation of membrane systems. This method immobilizes cells instantly by freezing, and fixatives dissolved in organic solvents, usually osmium tetroxide in acetone, penetrate the frozen tissues without deforming cellular structures. However, lipophilic compounds in sample tissues are extracted in acetone. Therefore, FS and CF are complementary methods to investigate ultrastructural membrane systems for lipid secretion.

In this study, we observed the hairy roots of *L. erythrorhizon* both by CF and FS in order to gain new insights about the membrane systems of plant cells secreting lipids. CF provided information on the localization of lipids in the cells of hairy roots. FS revealed the intact cellular ultrastructure concerning lipid secretion.

## Materials and Methods

### Plant materials

The hairy roots of *L. erythrorhizon* were generated as previously described (Tatsumi *et al*., 2020) and maintained in 1/2 Murashige-Skoog medium (Murashige and Skoog, 1962), where shikonin derivatives are less produced. For inducing shikonin production, the hairy roots were transferred into M9 medium without auxin and cytokinin and cultured on a rotary shaker (100 rpm) at 23 °C for 7 days in the dark. Stereo micrographs of the hairy roots of *L. erythrorhizon* that were cultured for 7 days in M9 medium are shown in Fig. 2. The root tips showing yellow color (Fig. 2b) and the mature parts in which red pigments were accumulated on the root hairs (Fig. 2c) were cut into about 3 mm length and were subjected to the fixation, embedding and microscopic observation as following.

**Fig. 2.**
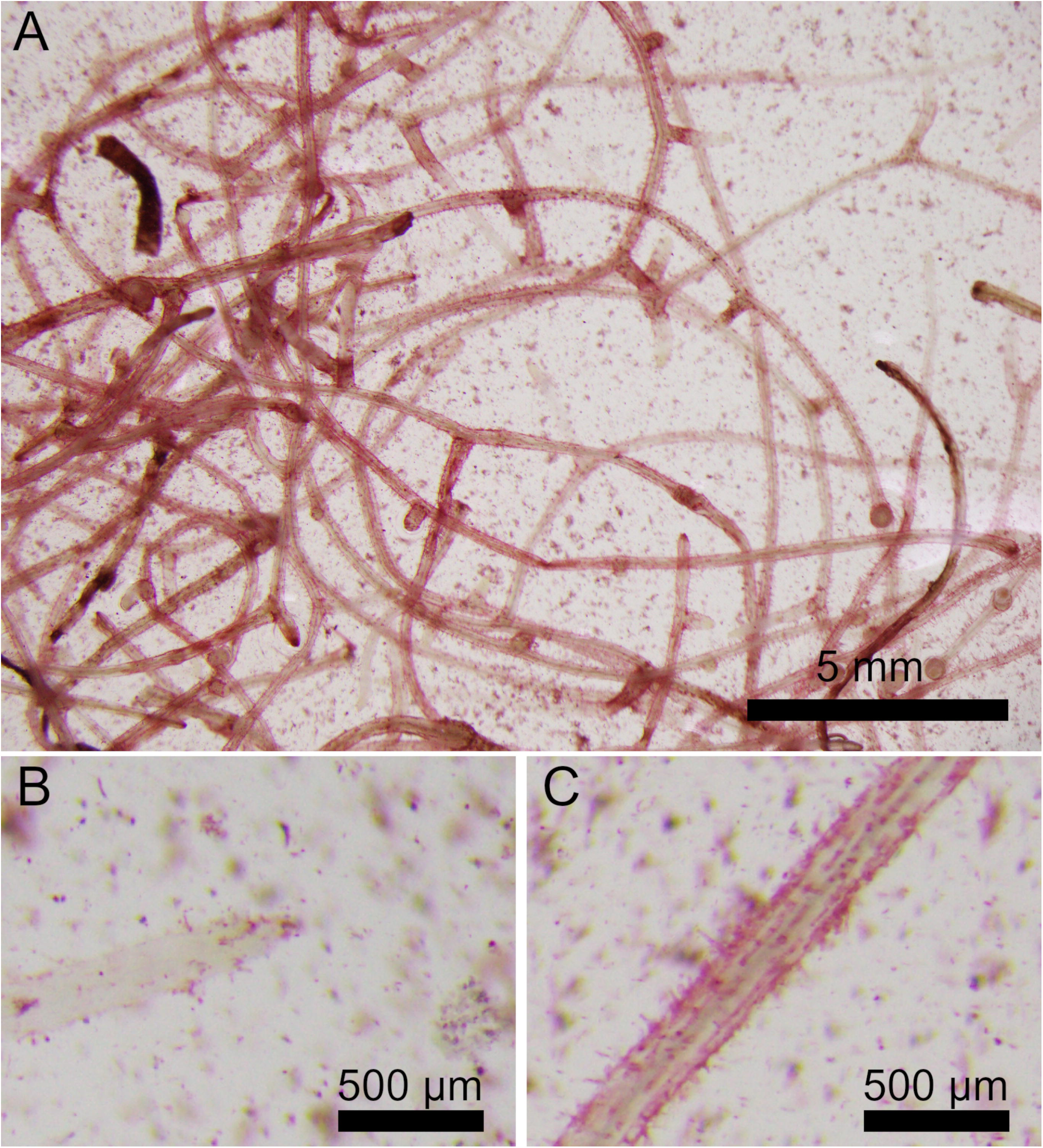
Stereoscopic micrographs of *L. erythrorhizon* hairy roots producing shikonin derivatives cultured in M9 medium for 7 days. (A) Low magnification image. High magnification image of the root tip (B), and mature parts (C).

### Spot tests

Spot tests were performed on a variety of compounds to see if they were retained by fixation and if each compound could be distinguished by color or shape. Ethanolic solutions of 10% (w/v) oleic acid, 1% (w/v) triolein, 1% ferulic (w/v) acid, and 1% (w/v) caffeic acid were prepared. An aliquot of these solutions (0.5 μL) was spotted onto filter paper. The spotting was repeated ten times for the compounds other than oleic acid. In addition, 1% (w/v) aqueous solutions of catechol and tannic acid were prepared. Pieces of filter paper were dipped into the solutions. The pieces of the filter paper were fixed and embedded in the same manner as the hairy root samples described in the following sections. However, instead of freezing in a high-pressure freezer, they were frozen by immersion in liquid nitrogen.

### Chemical fixation (CF)

To observe lipids in the hairy roots, improved CF was performed as described in the previous report (Kiyoto *et al*., 2022). The hairy roots were fixed with 3% (w/v) glutaraldehyde and 0.1% (w/v) malachite green oxalate in 50 mM Na-piperazine-N, N′-bis (2-ethane sulfonic acid) (Na-PIPES) buffer (pH 6.8) for 2 h at room temperature and overnight at 4°C. After washing five times with the buffer for 10 min each, the hairy roots were then post-fixed with 1% (w/v) osmium tetroxide in a solution containing 80 mM imidazole-HCl and 50 mM Na-PIPES buffer for 30 min at room temperature. After post-fixation, the sample was washed once with 80 mM imidazole-HCl in 50 mM Na-PIPES buffer (pH 6.8), twice with the Na-PIPES buffer, and three times with deionized water for 10 min each. After washing, samples were dehydrated with 30% and 50% ethanol, 10 min each. To enhance post-fixation with osmium tetroxide, the sample was transferred to 1% (w/v) *p*-phenylenediamine in 70% ethanol for 30 min. The sample was then dehydrated through an ethanol series (70%, 90% for 10 min each; 100% for 10 min, three times). After dehydration, samples were embedded in the Spurr Low Viscosity Embedding kit (Polysciences Inc., Warrington, PA, USA) according to the manufacturer’s instructions.

### Freeze-substitution after high-pressure freezing (HPF-FS)

Small pieces of the hairy root were excised with a surgical knife to a size that would fit into a Type B specimen carrier (2mm internal diameter, 300 µm depth; Leica Microsystems, Wetzlar, Germany) and placed on the carrier. The specimen carriers were filled with M9 medium in which sucrose concentration was adjusted to 0.2 M as cryoprotectant. The carrier was covered with another carrier and high-pressure frozen with a high-pressure freezer (Leica EM HPM100, Leica Microsystems, Wetzlar, Germany). The frozen samples were immersed in acetone solution containing 0.5% (w/v) osmium tetroxide, 40 mM imidazole and 1% (v/v) water which was pre-cooled in liquid nitrogen. FS was performed in a program freezer (CS-80CP, Scinics Corp., Tokyo, Japan). The sample was maintained at -80°C for 60 h; then warmed to -60°C at 2°C/h and held for 24 h; further warmed to -30°C at the same rate and held for another 24 h; then warmed to 0°C at 0.5°C/h and held for 2 h; finally, it was warmed to 20°C at 1°C/h. After FS, the samples were washed three times with acetone for 10 min each, once with 1% (w/v) *p*-phenylenediamine in ethanol for 30 min, three times with ethanol for 10 min each, and three times with propylene oxide for 10 min each. The samples were then stepwise embedded in the Spurr Low Viscosity Embedding kit according to the manufacturer’s instructions.

### Light microscopy

Transverse semithin sections (approx. 0.5 μm thick) were cut with an ultramicrotome from the samples fixed by CF. For the observation of the root tip, sections were cut from a region approx. 100 to 200 µm away from the tip. The sections were mounted in EUKITT® neo (O. Kindler & ORSAtec, Bobingen, Germany) without staining. The sections were observed under a light microscope (BX51, Olympus, Tokyo, Japan) equipped with a color CCD camera (DP72, Olympus).

### Transmission electron microscopy (TEM)

Transverse ultrathin sections (approx. 80 nm thick) were cut with an ultramicrotome and mounted on copper grids. The sections were stained with an aqueous solution of 1% (w/v) potassium permanganate and 0.1% (w/v) sodium citrate for 3 min at room temperature, washed with deionized water, and observed under a TEM (JEM1400, JEOL, Tokyo, Japan) at an accelerating voltage of 80 kV.

## Results

### Spot tests

Shikonin derivatives are secreted out of the cells together with triacylglycerol as matrix lipids (Tatsumi *et al*., 2023). Adding to shikonin derivatives, *L. erythrorhizon* also produces an appreciable amount of caffeic acid oligomers such as rosmarinic acid and lithospermic acid as well as monomers of simple phenols (Fukui *et al*., 1984; Yazaki *et al*., 1997; Yamamoto *et al*., 2000). Then, the following chemicals were used as standards in the spot tests; triolein, oleic acid, ferulic acid, caffeic acid, catechol, and tannic acid. Among the standard samples fixed by CF, triolein, oleic acid, and ferulic acid showed spherical shape (Figs. 3A, C, E). Caffeic acid showed amorphous shape (Fig. 3G). Catechol was not fixed (Fig. 3I). Tannic acid showed green-colored granular shape (Fig. 3K). Among the standard samples fixed by FS, triolein, oleic acid, caffeic acid, catechol, and tannic acid showed amorphous shape (Figs. 3B, D, H, J, L), while ferulic acid was not fixed (Fig. 3F).

**Fig. 3.**
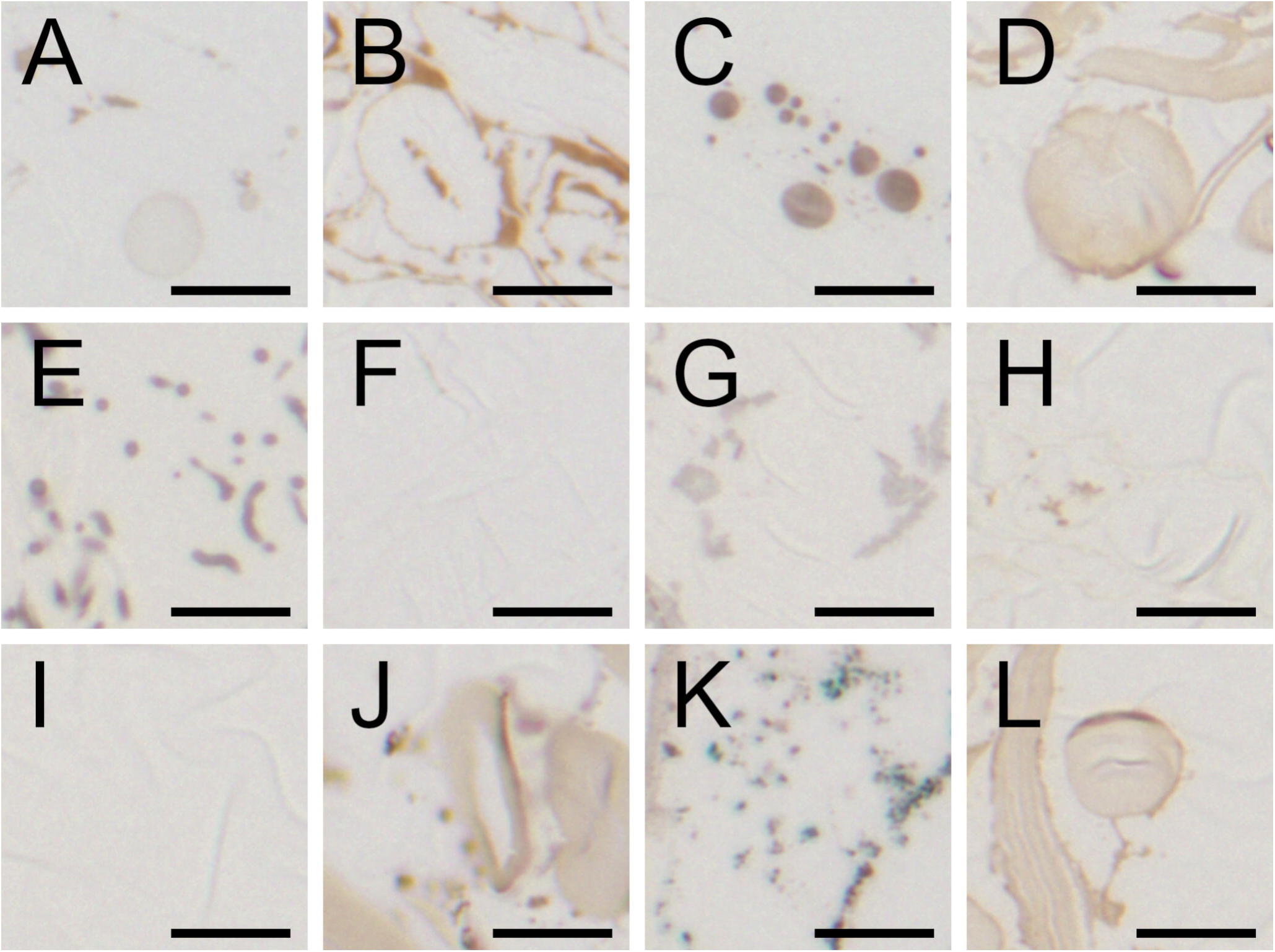
Spot tests on specific lipid standards. Light micrographs of filter paper fixed by CF (A, C, E, G, I, K) and the FS (B, D, F, H, J, L). (A, B) triolein, (C, D) oleic acid, (E, F) ferulic acid, (G, H) caffeic acid, (I, J) catechol, and (K, L) tannic acid, were spotted and fixed. The scale bars indicate 10 μm.

### Light microscopy

Light micrographs of cross sections of *L. erythrorhizon* hairy root fixed by CF are shown in Fig. 4. In the root tip, highly developed vacuoles were found in the epidermal cells while not in the cortical cells where cells were mostly filled with cytosol (Fig. 4A, B). In the mature part of hairy root, almost all cells had highly developed vacuoles (Fig. 4C). Black granules were observed mainly in the cytosol and sometimes in the vacuole in both the root tip and the mature part (Fig. 4B, D; black arrowheads). Green-colored materials were observed in the vacuole (Fig. 4B, D; arrows). Shikonin-containing granular structure attached were observed on cell surface (Fig. 4A, D; red arrowheads).

**Fig. 4.**
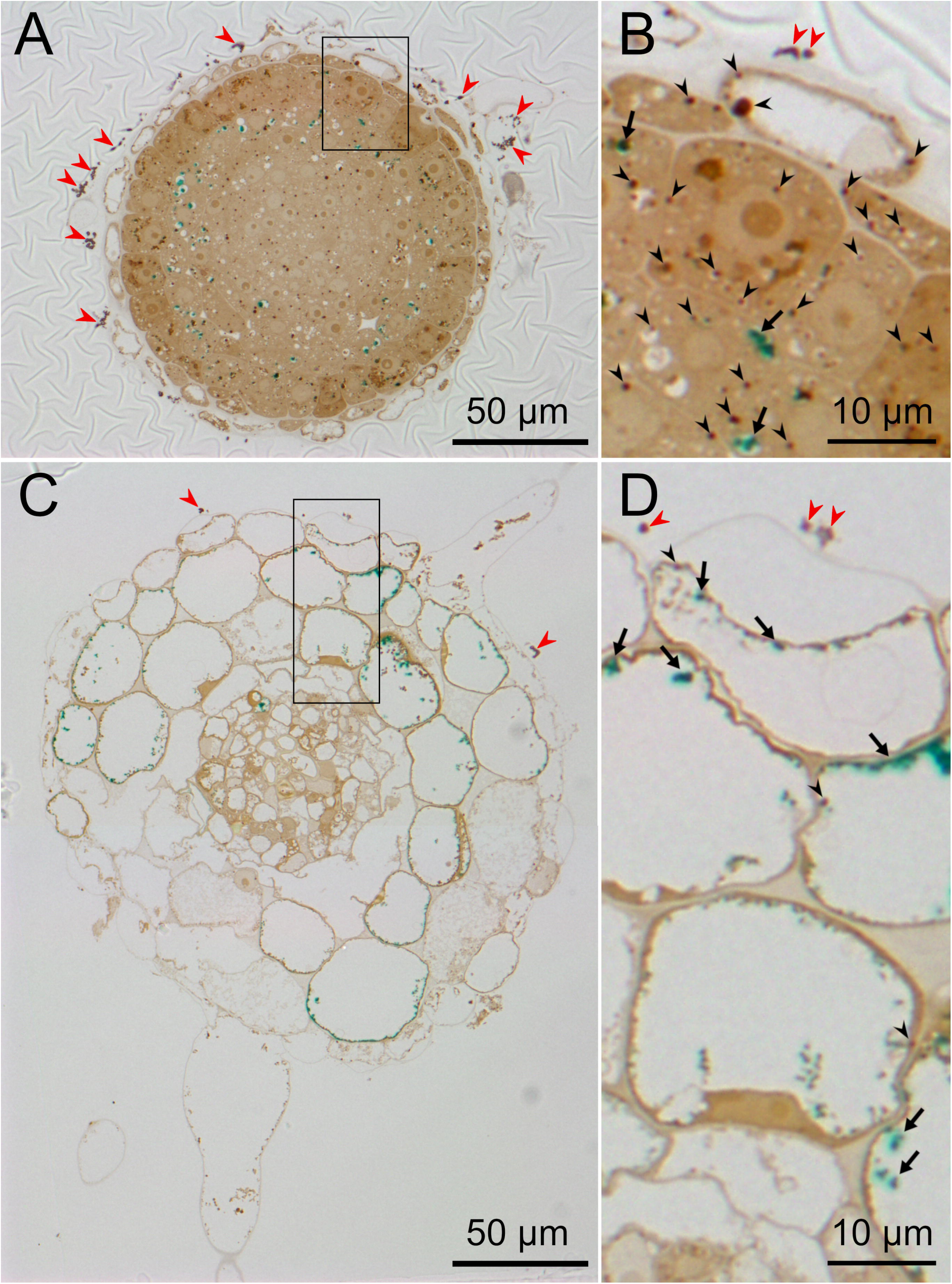
Light micrographs of *L. erythrorhizon* hairy roots producing shikonin derivatives fixed by CF. (A, B) Cross section at 100 to 200 μm length from the tip. (C, D) Cross section at the mature part of the root in which root hairs showed red color by the accumulation of shikonin granule. (B) Magnified image of the boxed region in (A). (D) Magnified image of the boxed region in (C). Black arrowheads indicate lipid droplets. Red arrowheads indicate shikonin granules. Arrows indicate green stained materials.

### Transmission electron microscopy

In the root tip fixed by CF, electron-dense lipid droplets were observed not only in the epidermal cells but also in the outer and inner cortical cells (Fig. 5A-I). In the vacuole of the epidermal cells, empty membrane structures were sometimes observed (Fig. 5B; arrowheads). The structure was attached to lipid droplets spanning the cytoplasm and vacuole (Fig. 5C; arrowheads). Small electron-dense grains were sometimes observed both inside and outside of epidermal cell (Fig. 4D), and also in the epidermal cell wall (Fig. 5E; arrowheads). Lipid droplets enveloped in the multivesicular bodies (MVBs) were observed both in the outer and inner cortical cells (Fig. 5F, H; arrowheads). In addition, the invagination of plasma membrane was observed (Fig. 5F; arrow), sometimes with lipid droplet (Fig. 5H; arrow). A lipid droplet spanning the cytoplasm and MVBs was observed (Fig. 5I; arrowheads).

**Fig. 5.**
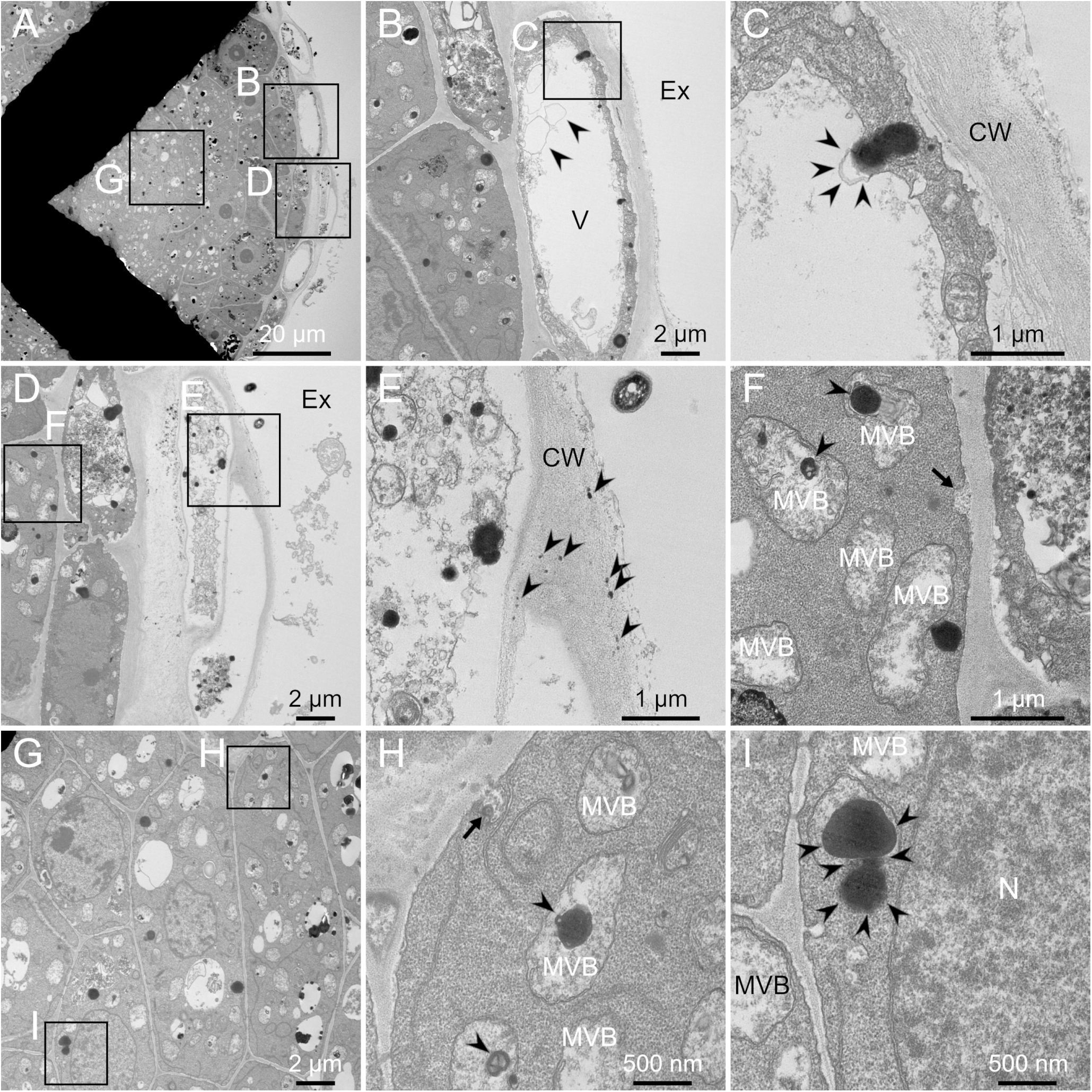
TEM images of the cross section cut from the area about 100 to 200 μm length from the tip of *L. erythrorhizon* hairy root fixed by CF. The sections were stained with potassium permanganate. (A) Low magnification image. (B, D, G) Magnified image of the boxed region in (A). (B) Arrowheads indicate the empty membrane structure in the vacuole. (C) Magnified image of the boxed region in (B). Arrowheads indicate the empty membrane structure in the vacuole. (E, F) Magnified image of the boxed region in (D). (E) Arrowheads indicate electron-dense grains in the cell wall. (F) Arrowheads indicate electron-dense bodies surrounded by MVBs. The arrow indicates an MVB fusing with the plasma membrane. (H, I) Magnified image of the boxed region in (G). (H) Arrowheads indicate electron-dense bodies in MVBs. The arrow indicates an electron-dense body in the MVB fusing with the plasma membrane. (I) Arrowheads indicate a lipid droplet spanning the cytosol and MVBs. CW, cell wall; Ex, extracellular spaces; MVB, multivesicular body; N, nucleus; V, vacuole.

In the root tip fixed by FS, vacuoles in the epidermal cells were filled with dispersed materials or electron-dense materials (Fig. 6A and B, respectively). Vacuoles filled with electron-dense materials were also observed in the cortical cells and sometimes branched (Fig. 6C; arrowheads). MVBs contacted with plastids (Fig. 6D; arrowheads) or with lipid droplets (Fig. 6E; arrowheads) were sometimes observed. The invagination of plasma membrane was observed and showed similar texture to MVBs (Fig. 6F; arrowheads).

**Fig. 6.**
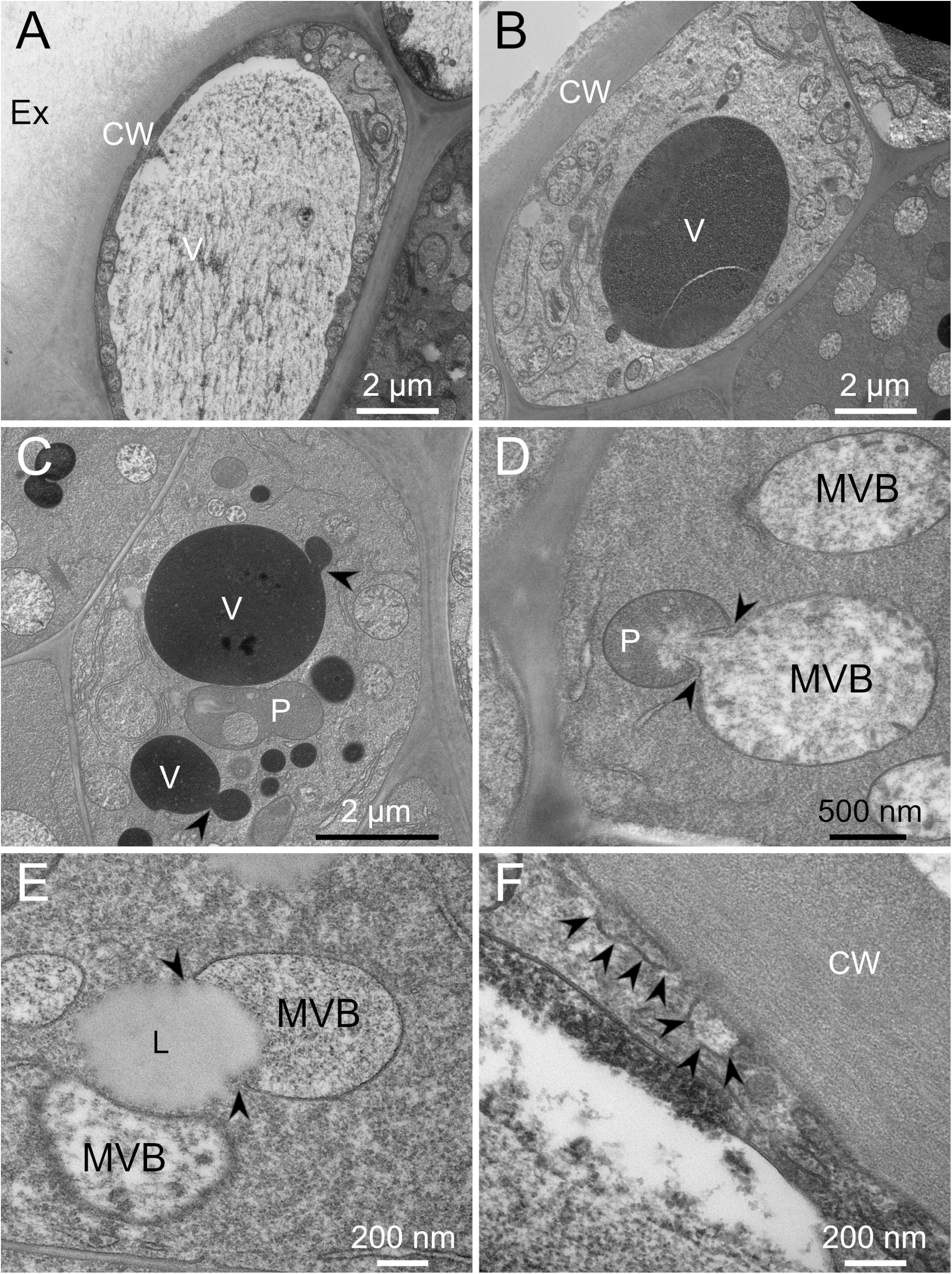
TEM images of the cross section cut from the area about 100 to 200 μm length from the tip of *L. erythrorhizon* hairy root fixed by FS. The sections were stained with potassium permanganate. (A, B) Epidermal cells. (C, D, E, F) Cortical cells. (A) The vacuole is filled with dispersed materials. (B) The vacuole is filled with electron-dense materials. (C) Arrowheads indicate branches of vacuoles filled with electron-dense materials (D) Arrowheads indicate an MVB connecting with a plastid. (E) Arrowheads indicate an MVB taking up a lipid droplet. (F) Arrowheads indicate the invagination of plasma membrane considered to be MVBs fusing with plasma membrane. CW, cell wall; Ex, extracellular spaces; L, lipid droplet; MVB, multivesicular body; P, plastid; V, vacuole.

TEM images of the mature parts fixed by CF were shown in Fig. 7A-I. Electron-dense bodies not only existed in MVBs but also straddle between two MVBs (Fig. 7B; arrowhead). Electron-dense bodies were also observed in the invagination of plasma membrane (Fig. 7B; arrows) and in the cell wall (Fig. 7C, H; arrows). Empty membrane structures in the vacuoles sometimes fused with the MVBs (Fig. 7C; arrowhead), or cytosols (Fig. 7H, I; arrowheads). Shikonin granules secreted to the extracellular spaces showed variation in the textures: Some showed vacuole-like structures (Fig. 7D), and others were filled with electron-dense materials (Fig. 7E). Developed endoplasmic reticulum (ER) was observed both in the epidermal and cortical cells (Fig. 7E, G).

**Fig. 7.**
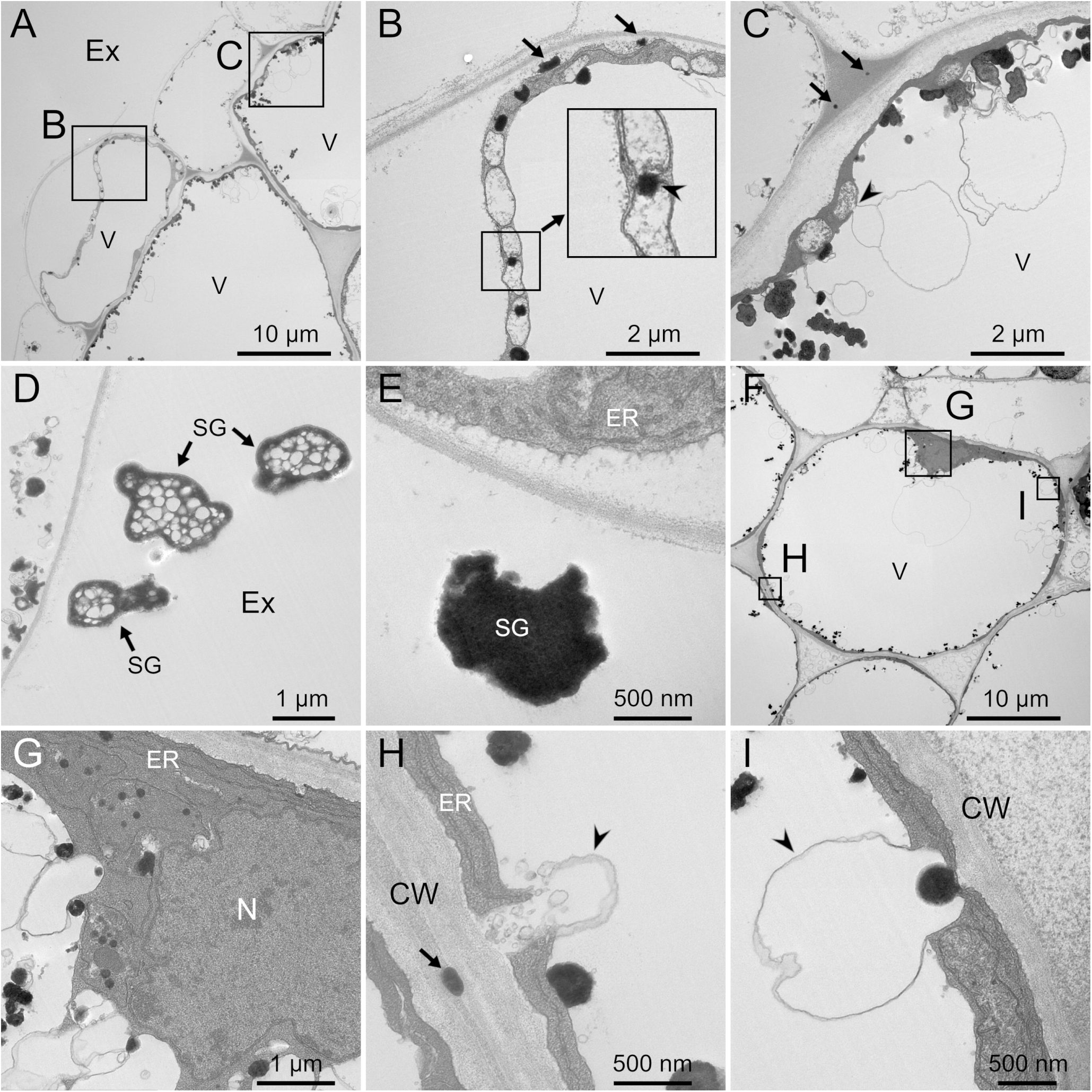
TEM images of the cross section at the mature part of *L. erythrorhizon* hairy root fixed by CF. (A, B, C, D, E) Epidermal cells. (F, G, H, I) A cortical cell. (B, C) Magnified image of the boxed region in (A). (B) Arrowhead indicates lipid droplet straddling two MVBs. Arrows indicate lipid droplets in the invagination of plasma membrane considered to be MVBs fusing plasma membrane. (C) Arrowhead indicates connection site of an MVB and an empty membrane structure in the vacuole. Arrows indicate lipids passing through the cell wall. (D) Shikonin granules show vacuole-like structures. (E) Shikonin granule is filled with electron-dense materials. (F) Low magnification image. (G, H, I) Magnified image of the boxed region in (F). (G) ER is developed. Lipid droplets were often observed. (H) Arrowhead indicates the empty membrane structure fusing with cytosol. Arrow indicates lipids in the cell wall. (I) Arrowhead indicates the empty membrane structure fusing with cytosol. CW, cell wall; Ex, extracellular spaces; ER, endoplasmic reticulum; N, nucleus; SG, shikonin granule; V, vacuole.

TEM images of the mature parts fixed by FS were shown in Fig. 8A-F. Vacuoles were filled in dispersed materials in the epidermal (Fig. 8A, B, C) and cortical cells (Fig. 8D, E, F). Shikonin granules secreted to extracellular spaces showed various textures: Some were empty, some were filled with electron-dense grains, and others were filled with electron-dense materials (Fig. 8A, B, C; arrowheads). Plasma membranes were smoothly fitted to the cell wall in the cortical cells (Fig. 8D), but the invagination of plasma membrane was sometimes observed and showed similar texture to MVBs (Fig. 8F; arrowheads). A connection between two MVBs was observed (Fig. 8E; arrowhead).

**Fig. 8.**
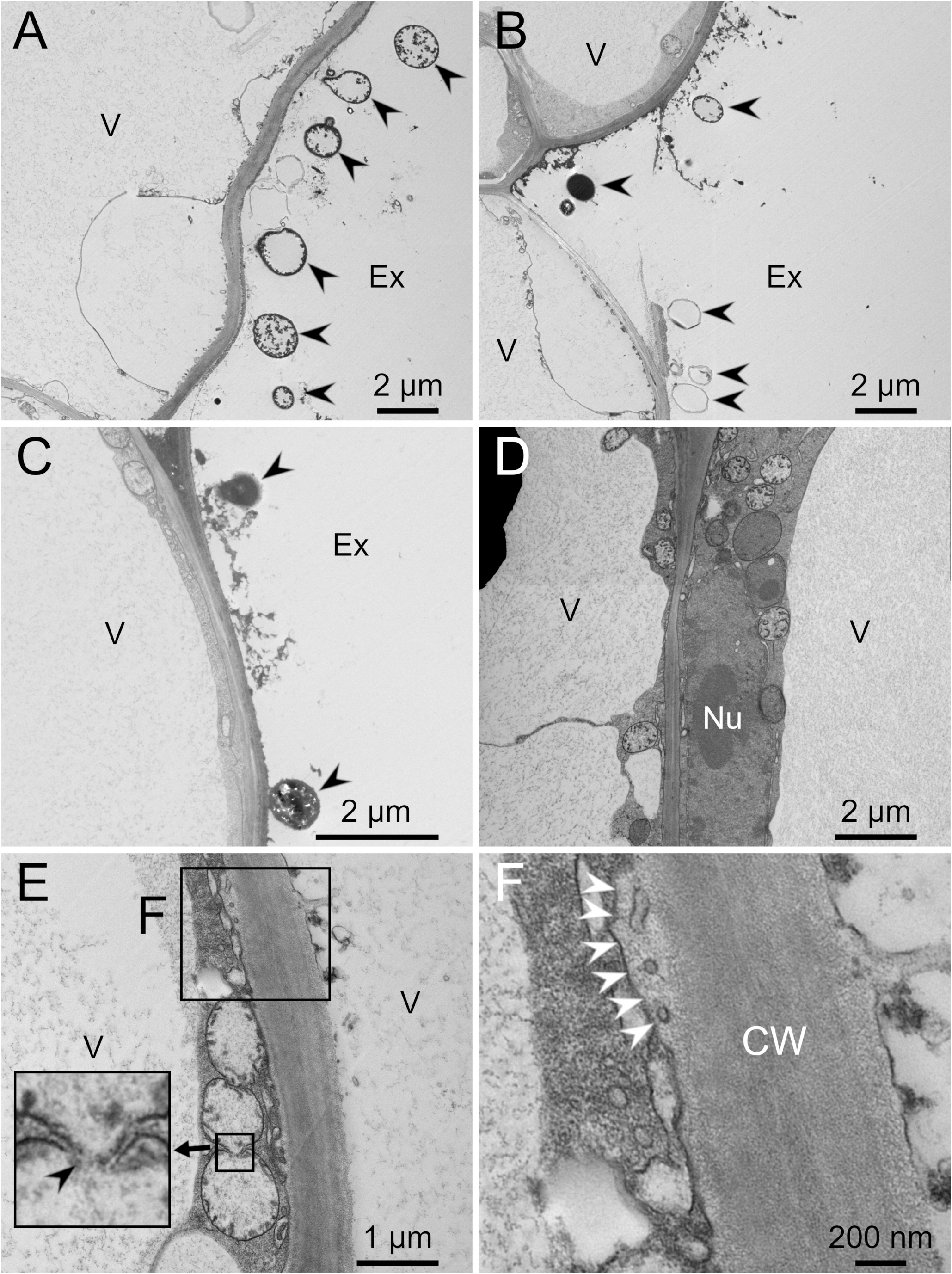
TEM images of the cross section at the mature part of *L. erythrorhizon* hairy root fixed by FS. The sections were stained with potassium permanganate. (A, B, C) Epidermal cells. Arrowheads indicate shikonin granules secreted to the extracellular space. (D) Cortical cells. Vacuoles were filled with dispersed materials. (E) Cortical cells. Arrowhead indicates connection site of two MVBs. (F) Magnified image of the boxed region in (E). Arrowheads indicate the invagination of plasma membrane considered to be MVBs fusing with plasma membrane. CW, cell wall; Ex, extracellular spaces; Nu, nucleous; V, vacuole.

## Discussion

### Effects and mechanisms of the fixatives

Osmium tetroxide acts as both a fixative and an electron-staining reagent and is commonly used in both CF and in FS. This reagent reacts with the side chain of cysteine, methionine and tryptophan residues in proteins (Bland *et al*., 1971; Deetz and Behrman, 1980), as well as with carbon double bonds (Korn, 1967; Bland *et al*., 1971), and 1,2-dihydroxybenzene (catechol) structures (Bland *et al*., 1971; Nielson and Griffith, 1978). It means that proteins, phospholipid membranes, and several lipophilic metabolites which have carbon double bonds or catechol structure are fixed and electronically stained by osmium tetroxide. Imidazole and *p*-phenylenediamine enhance electron staining and fixation of membrane and lipids by osmium tetroxide (Angermüller and Fahimi 1984; Ledingham and Simpson 1972; Nakao *et al*. 1992; Shikata *et al*., 1989). Tatsumi *et al*. (2023) reported that formation of shikonin-containing lipid droplets requires the presence of unsaturated triacylglycerol. In other words, the lipid droplets in *L. erythrorhizon* cells contain not only shikonin derivatives but also other lipophilic compounds such as unsaturated triacylglycerol. In addition, plant cells produce osmiophilic compounds such as phenylpropanoids and caffeic acid derivatives produced by *L. erythrorhizon* are representative. The osmiophilic metabolites in plant cells include both lipophilic and hydrophilic compounds. Therefore, each compound is expected to show different fixation behavior depending on whether it is lipophilic or hydrophilic and whether it is fixed by CF or FS. To investigate their fixation behavior in CF and FS, several osmiophilic standard materials were subjected to spot tests.

In the spot test, triolein and oleic acid showed a spherical shape by CF but showed an amorphous shape by FS (Fig. 3A, B, C, D). This is likely because these hydrophobic compounds retained their spherical shape during the aqueous CF, whereas they were deformed by dissolution in acetone before being fixed by osmium tetroxide in FS. Ferulic acid was fixed by CF but extracted by FS, while caffeic acid was slightly fixed by FS (Fig. 3E, F, H). It indicates that a carbon double bond conjugated with an aromatic ring is less reactive to osmium tetroxide than one without conjugation or a catechol structure. Therefore, ferulic acid was completely extracted from the filter paper before fixation. In any case, the lipophilic compounds cannot retain their original position and shape due to dissolution in acetone when fixed by FS. As the result, lipid droplets showed lower electron density in the samples fixed by FS than those by CF (Figs. 5, 6E, 7B, G). Catechol was fixed by FS but not by CF (Fig. 3I, J). Because the compound is water-soluble, it would have been extracted at the stage of the pre-fixation in an aqueous system. Tannic acid is also highly water-soluble, but it was fixed by CF and showed green color probably because of malachite green (Fig. 3K). We cannot explain the detailed mechanism, but these results indicate that the CF procedure in this research can distinguish tannic acid from other compounds by the green color.

### Ultrastructural membrane systems for transport and secretion of shikonin derivatives

*L. erythrorhizon* hairy roots producing shikonin derivatives show red color mainly in the mature part by accumulation of the pigments (Fig. 2C). Therefore, past TEM observation of the hairy roots was performed on mature parts (Tatsumi *et al*., 2016). However, the same paper also reported that the hairy roots accumulate shikonin derivatives on the root tip by the inhibitor treatment of membrane trafficking, such as cytochalasin D and brefeldin A. It indicates the ability to produce shikonin derivatives in the root tips of hairy roots. Cells in the root tip are rich in cytosol, while those in the mature part have developed vacuole, resulting in a reduced cytosolic area in the latter (Fig. 4). Therefore, the root tip of *L. erythrorhizon* hairy roots is a good sample for the investigation of lipid production and secretion in cytosol area.

In the vacuoles of the hairy roots, the green stained materials were observed (Fig. 4). They were also observed in the cells cultured with M9 medium (Kiyoto *et al*., 2022). As mentioned above, the CF procedure in this research can stain tannic acid green. The results indicate that shikonin-producing *L. erythrorhizon* produces tannin-like polyphenolic compounds in the vacuole. Here, the production of lithospermic acid B, a tetramer of caffeic acid, is similarly induced in M9 medium as that of shikonin derivatives (Yamamoto *et al*., 2020). This indicates that the green-stained materials may be lithospermic acid B and/or related compound like rosmarinic acid that is a dimer of caffeic acid (Fukui *et al*., 1984). The localization of the metabolites in vacuole corresponds to the past report that lithospermic acid B may be accumulated in the vacuole (Yazaki *et al*., 2017).

The vacuoles of several epidermal and cortical cells in the root tip fixed by FS were filled with electron-dense materials though those fixed by CF were not (Figs. 5A, B, D, 6B, C). Here, as mentioned above, catechol was fixed by FS but not by CF (Fig. 3I, J). These results indicate that the vacuoles of several cells in the root tip are filled with water-soluble osmiophilic compounds similar to catechol. The precursors of above mentioned caffeic acid oligomers are phenylpropanoid monomers, and these are putative candidates of the electron-dense materials.

ER plays an important role in the biosynthesis and the secretion of lipids including shikonin, as reported previously (Tsukada and Tabata, 1984; Tabata, 1996; Tatsumi *et al*., 2016; Kiyoto *et al*., 2022). These observations are consistent with the understandings that lipids such as triacylglycerol are synthesized in ER and that these lipid droplets are budding out from the ER (Jacquemyn *et al*., 2017; Choudhary and Schneiter, 2020). Here, shikonin derivatives accumulates mainly in the epidermal cells in the dark (Yazaki *et al*., 1999; Yazaki, 2017). However, lipid droplets and developed ER were often observed both in the epidermal and cortical cells (Figs. 5C, H, 6A, B, 7E, G). It may indicate that not only epidermal cells but also cortical cells produce lipids, which may include the precursors of shikonin.

Empty membrane structures were sometimes observed in developed vacuoles both in epidermal and cortical cells in the samples fixed by CF (Figs. 5B, C, 7C, F, H, I). In some cases, they attached to or surrounded electron-dense bodies (Figs. 5C, 7I). In other cases, they fused with MVBs (Fig. 7C; arrowhead). These observations imply that the empty membrane structures are the machinery for transporting compounds between vacuoles and cytosols/MVBs.

Shikonin granules observed in the extracellular spaces showed variable textures (Figs. 7D, E, 8A, B, C). It is consistent with the variable fixation behaviors of osmiophilic compounds (Fig. 3). The observation results also implies that each shikonin granule has significantly different composition. To identify the compounds in the electron-dense materials in such as vacuole, lipid droplets, and shikonin granules, further investigation is required.

For secretion of lipids to extracellular spaces, there must be the transport pathway in the cell walls. The electron-dense grains which seems to be during passage through the cell walls have variability in the size (Figs. 5E, 7C, H). To know the shape and size of the pathway for lipids in cell walls, three-dimensional observation is required such as FIB-SEM tomography (Kuo, 2014).

In the samples fixed by CF, MVBs surrounded lipid droplets (Figs. 5F, H, I; arrowheads, 7B). In addition, the invagination of plasma membrane showed similar textures to MVBs (Fig. 5F; arrows) and sometimes surrounded electron-dense bodies (Figs. 5H, 7B; arrows). The invagination of plasma membrane showing similar textures to MVBs were also observed in the samples fixed by FS (Figs. 6F, 8F; arrowheads), indicating that they are not artifacts such as bacterial mesosome, but are membrane fusion of MVBs and plasma membranes. The membrane fusion surrounding electron-dense body was observed not only in the direction to the extracellular space (Fig. 7B; arrows) but also in the opposite side (Fig. 5H; arrow). This suggests that MVBs may be involved in both the exocytosis and endocytosis of shikonin derivative-containing lipids. De Bellis *et al*. (2022) reported that the presence of extracellular vesicles, MVBs fusing with the plasma membrane, correlates with root suberization of *Arabidopsis thaliana* and hypothesized that extracellular vesicles are involved in transport and secretion of suberin monomer. Suberin monomers, fatty acids and phenylpropanoids, are also lipophilic compounds as with shikonin derivatives, and are secreted to extracellular spaces for cell wall suberization. There may be similar machinery for their transport and secretion.

The aromatic precursor of shikonin, *p*-hydroxybenzoate, is derived from shikimate pathway localized in plastids, which is responsible for the very upstream biosynthetic steps (Yazaki, 2017). Recently, the involvement of plastids in shikonin biosynthesis has also been suggested by the genome editing experiments of polyphenol oxidase (PPO), which is localized in plastids (Nakanishi *et al*., 2025, Preprint). This biosynthetic reaction catalyzed by PPO is proposed to be very downstream of the shikonin biosynthesis such as naphthalene ring formation. Besides, the key intermediate geranyl pyrophosphate is produced in the cytosol via the mevalonate pathway (Ueoka *et al*., 2020). Furthermore, the enzyme LePGT, which couples *p*-hydroxybenzoate with geranyl pyrophosphate, is localized to the ER (Tatsumi *et al*., 2023). This localization of multiple biosynthetic enzymes suggests a complex interplay between plastids and other cellular compartments in shikonin biosynthesis. The observation of an MVB in contact with a plastid (Fig. 6D; arrowheads) indicates that MVBs may also be involved in the transport of the materials for the biosynthesis of shikonin. These observations may share a common phenomenon observed in cannabinoids in hemp, which are also lipophilic metabolites and secreted out of the cells (Livingston *et al*., 2022).

Electron-dense bodies straddling between two MVBs (Fig. 7B; arrowhead) may mean the migration of lipids between the two MVBs. The observation of two connected MVBs in the samples fixed by FS (Fig. 8E; arrowhead) supports this idea. The idea also indicates that each MVB plays a different role such as in exocytosis, endocytosis, intracellular lipid transport, and lipid uptake from the cytosol, vacuoles or other organelles.

In conclusion, TEM observation of *L. erythrorhizon* hairy roots fixed by CF found that MVBs are involved in biosynthesis and secretion of lipids considered to contain shikonin derivatives and that both the epidermal and cortical cells produce the lipids. However, many questions remain unclear. What is the detailed role of the empty membrane structures observed in the vacuoles? How lipids pass through the cell wall? What are the detailed compositions of the lipid droplets observed in the CF and the electron-dense vacuoles observed in the FS? Further investigations are required for unraveling these questions.

## Abbreviations

CF: Chemical fixation
ER: Endoplasmic reticulum
FS: Freeze-substitution
MVB: Multivesicular body
TEM: Transmission electron microscopy

## Acknowledgments

We are grateful to Dr. Koichiro Shimomura (Toyo University, Japan) and Dr. Noboru Onishi (Kirin Holdings Company Limited, Japan) for sharing axenic shoots of *L. erythrorhizon*. We also thank Dr. Masahiro Mii and Dr. Tomoko Igawa (Chiba University, Japan) for sharing *R. radiobacter* strain A13.

## Author contributions

KY, SK and TA: conceptualization; SK: methodology; SK: investigation; TI: resources; SK: data curation; SK: writing - original draft; TA, TI and KY: writing - review & editing; SK: visualization; KY: supervision; KY: funding acquisition

## Data availability statement

The raw TEM images and other primary data supporting the findings of this study are not publicly available, but are available from the corresponding author on reasonable request.

## References

Angermüller S, Fahimi HD. 1982. Imidazole-buffered osmium tetroxide: an excellent stain for visualization of lipids in transmission electron microscopy. The Histochemical Journal 14, 823–835.

Bland DE, Foster RC, Logan AF. 1971. The mechanism of permanganate and osmium tetroxide fixation and the distribution of lignin in the cell wall of *Pinus radiata*. Holzforschung 25, 137–143.

Choudhary V, Schneiter R. 2020. Lipid droplet biogenesis from specialized ER subdomains. Microbial Cell 7, 218–221.

Clews AC, Ulch BA, Jesionowska M, Hong J, Mullen RT, Xu Y. 2024. Variety of Plant Oils, Species-Specific Lipid Biosynthesis. Plant and Cell Physiology 65, 845–862.

De Bellis D, Kalmbach L, Marhavy P, Daraspe J, Geldner N, Barberon M. 2022. Extracellular vesiculo-tubular structures associated with suberin deposition in plant cell walls. Nature Communications 13, 1489.

Deetz JS, Behrman EJ. 1981. Reaction of osmium reagents with amino acids and proteins. International Journal of Peptide and Protein Research 17, 495–500.

Ebersold HR, Cordier JL, Ltithy P. 1981. Bacterial mesosomes, method dependent artifacts. Archives of Microbiology 130, 19–22.

Fu X, Han H, Wang Y, Liu H, Liu P, Li L, Zhao J, Sun X, Tang K. 2024. AaABCG20 transporter involved in cutin and wax secretion affects the initiation and development of glandular trichomes in *Artemisia annua*. Plant Science 339, 111959.

Fujita Y, Hara Y, Suga C, Morimoto T. 1981. Production of shikonin derivatives by cell suspension cultures of *Lithospermum Erythrorhizon* II. A new medium for the production of shikonin derivatives. Plant Cell Reports 1, 61–63.

Fukui H, Yazaki K, Tabata M. 1984. Two phenolic acids from *Lithospermum erythrorhizon* cell suspension cultures. Phytochemistry 23, 2398–2399.

Guo C, He J, Song X, Tan L, Wang M, Jiang P, Li Y, Cao Z, Peng C. 2019. Pharmacological properties and derivatives of shikonin-a review in recent years. Pharmacological Research 149, 104463.

Hagström L, Bahr GF. 1960. Penetration of osmium tetroxide with different fixation vehicles. Histchemie 2, 1–4.

Ichino T, Maeda K, Hara-Nishimura I, Shimada T. 2020. Arabidopsis ECHIDNA protein is involved in seed coloration, protein trafficking to vacuoles, and vacuolar biogenesis. Journal of Experimental Botany 71, 3999–4009.

Ichino T, Yazaki K. 2022. Modes of secretion of plant lipophilic metabolites via ABCG transporter-dependent transport and vesicle mediated trafficking. Current Opinion in Plant Biology 66, 102184.

Jacquemyn J, Cascalho A, Goodchild RE. 2017. The ins and outs of endoplasmic reticulum-controlled lipid biosynthesis. EMBO reports 11, 1905–1921.

Kiyoto S, Ichino T, Awano T, Yazaki K. 2022. Improved chemical fixation of lipid-secreting plant cells for transmission electron microscopy. Microscopy 71, 206–213.

Korn ED. 1967. A chromatographic and spectrophotometric study of the products of the reaction of osmium tetroxide with unsaturated lipids. Journal of Cell Biology 34, 627–638.

Kuo J. 2014. Electron microscopy methods and protocols. Humana Press: Totowa.

Ledingham JM, Simpson FO. 1972. The use of *p*-phenylenediamine in the block to enhance osmium staining for electron microscopy. Stain Technology 47, 239–243.

Li H, Matsuda H, Tsuboyama A, Munakata R, Sugiyama A, Yazaki K. 2022. Inventory of ATP-binding cassette proteins in *Lithospermum erythrorhizon* as a model plant producing divergent secondary metabolites. DNA Research 29, 1–12.

Livingston SJ, Rensing KH, Page JE, Samuels AL. 2022. A polarized supercell produces specialized metabolites in cannabis trichomes. Current Biology 32, 4040–4047.

McFarlane HE, Shin JJH, Bird DA, Samuels AL. 2010. Arabidopsis ABCG transporters, which are required for export of diverse cuticular lipids, dimerize in different combinations. Plant Cell 22, 3066–3075.

McFarlane HE, Watanabe Y, Yang W, Huang Y, Ohlrogge J, Samuels AL. 2014. Golgi-and trans-Golgi network-mediated vesicle trafficking is required for wax secretion from epidermal cells. Plant Physiology 164, 1250–1260.

Murashige T, Skoog F. 1962. A revised medium for rapid growth and bio assays with tobacco tissue cultures. Physiologia Plantarum 15, 473–497.

Nakanishi K, Takano Y, Yamamoto K, et al. CRISPR/Cas9-mediated genome editing reveals the involvement of a polyphenol oxidase in the shikonin-specific biosynthesis in *Lithospermum erythrorhizon*. bioRxiv doi: 10.1101/2025.07.30.667576 [Preprint]

Nakao I, Okada H, Senzaki H, Nishimura R, Fujita N, Shikata N, Morii S. 1992. A mixture of paraphenylenediamine and imidazole: its effect on the extraction of lipid droplets during electron microscopy staining. Biotechnic & Histochemistry 67, 219–223.

Nanninga N. 1968. Structural features of mesosomes (chondrioids) of *bacillus subtilis* after freeze-etching. Journal of Cell Biology 39, 251–263.

Nanninga N. 1969. Preservation of the ultrastructure of *bacillus subtilis* by chemical fixation as verified by freeze-etching. Journal of Cell Biology 42, 733–744.

Nanninga N. 1971. The mesosome of bacillus subtilis as affected by chemical and physical fixation. Journal of Cell Biology 48, 219–224.

Nielson AJ, Griffith WP. 1978. Tissue fixation and staining with osmium tetroxide: the role of phenolic compounds. Journal of Histochemistry & Cytochemistry 26, 138–140.

Pighin JA, Zheng H, Balakshin LJ, Goodman IP, Western TL, Jetter R, Kunst L, Samuels AL. 2004. Plant cuticular lipid export requires an ABC transporter. Science 306, 702–704.

Shikata N, Izuno Y, Nakao I, Morii S. 1989. Enhanced contrast of biomembranes with paraphenylenediamine- or imidazole-containing acetone after freeze-substitution fixation. Acta Histochemica et Cytochemica 22, 735–736.

Shimomura K, Sudo H, Saga H, Kamada H. 1991. Shikonin production and secretion by hairy root cultures of *Lithospermum erythrorhizon*. Plant Cell Reports 10, 282–285.

Tabata M, Mizukami H, Hiraoka N, and Konoshima M. 1974. Pigment formation in callus cultures of *Lithospermum erythrorhizon*. Phytochemistry 13, 927–932.

Tatsumi K, Yano M, Kaminade K, Sugiyama A, Sato M, Toyooka K, Aoyama T, Fumihiko Sato F, Yazaki K. 2016. Characterization of shikonin derivative secretion in *Lithospermum erythrorhizon* hairy roots as a model of lipid-soluble metabolite secretion from plants. Frontiers in Plant Science 7, 1066.

Tatsumi K, Ichino T, Isaka N, et al. 2023. Excretion of triacylglycerol as a matrix lipid facilitating apoplastic accumulation of a lipophilic metabolite shikonin., Journal of Experimental Botany 74, 104–117.

Tabata M. 1996. The mechanism of shikonin biosynthesis in cell cultures. Plant tissue culture letters 13, 117–125.

Tabata M, Mizukami H, Hiraoka N, Konoshima M. 1974. Pigment formation in callus cultures of *Lithospermum erythrorhizon*. Phytochemistry 13, 927–932.

Tsukada M, Tabata M. 1984. Intracellular localization and secretion of naphthoquinone pigments in cell cultures of *Lithospermum erythrorhizon*. Planta Medica 50, 338–341.

Ueoka H, Sasaki K, Miyawaki T, et al. 2020. A cytosol-localized geranyl diphosphate synthase from *Lithospermum erythrorhizon* and its molecular evolution. Plant Physiology 182, 1933–1945.

Yamamoto H, Tsukahara M, Yamano Y, Wada A, Yazaki K. 2020. Alcohol dehydrogenase activity converts 3’-hydroxygeranylhydroquinone to an aldehyde intermediate for shikonin and benzoquinone derivatives in *Lithospermum erythrorhizon*. Plant and Cell Physiology 61, 1798–1806.

Yamamoto H, Yazaki K, Inoue K. 2000. Simultaneous analysis of shikimate-derived secondary matabolites in *Lithospermum erythrorhizon* cell suspension cultures by high-performance liquid chromatography. Journal of Chromatography A 738, 3–15.

Yazaki K. 2017. *Lithospermum erythrorhizon* cell cultures, Present and future aspects. Plant Biotechnology 34, 131–142.

Yazaki K, Fukui H, Nishikawa Y, Tabata M. 1997. Measurement of phenolic compounds and their effect on shikonin production in *Lithospermum* cultured cells. Bioscience, Biotechnology, and Biochemistry 61, 1674–1678.

Yazaki K, Matsuoka H, Ujihara T, Sato F. 1999. Shikonin biosynthesis in *Lithospermum erythrorhizon*, Light-induced negative regulation of secondary metabolism. Plant Biotechnology 16, 335–342.

